# SPOTlight: Seeded NMF regression to Deconvolute Spatial Transcriptomics Spots with Single-Cell Transcriptomes

**DOI:** 10.1101/2020.06.03.131334

**Authors:** Marc Elosua, Paula Nieto, Elisabetta Mereu, Ivo Gut, Holger Heyn

## Abstract

The integration of orthogonal data modalities greatly supports the interpretation of transcriptomic landscapes in complex tissues. In particular, spatially resolved gene expression profiles are key to understand tissue organization and function. However, spatial transcriptomics (ST) profiling techniques lack single-cell resolution and require a combination with single-cell RNA sequencing (scRNA-seq) information to deconvolute the spatially indexed datasets. Leveraging the strengths of both data types, we developed SPOTlight, a computational tool that enables the integration of ST with scRNA-seq data to infer the location of cell types and states within a complex tissue. SPOTlight is centered around a seeded non-negative matrix factorization (NMF) regression, initialized using cell-type marker genes, and non-negative least squares (NNLS) to subsequently deconvolute ST capture locations (spots). Using synthetic spots, simulating varying reference quantities and qualities, we confirmed high prediction accuracy also with shallowly sequenced or small-sized scRNA-seq reference datasets. We trained the NMF regression model with sample-matched or external datasets, resulting in accurate and sensitive spatial predictions. SPOTlight deconvolution of the mouse brain correctly mapped subtle neuronal cell states of the cortical layers and the defined architecture of the hippocampus. In human pancreatic cancer, we successfully segmented patient sections into healthy and cancerous areas, and further fine-mapped normal and neoplastic cell states. Trained on an external pancreatic tumor immune reference, we charted the localization of clinical-relevant and tumor-specific immune cell states. Using SPOTlight to detect regional enrichment of immune cells and their co-localization with tumor and adjacent stroma provides an illustrative example in its flexible application spectrum and future potential in digital pathology.

## Introduction

Spatially resolved transcriptomics is key in advancing our understanding of tissue architectures. Unveiling the spatial disposition of cells enables researchers to determine cell-cell interactions and tissue reconstruction for a better knowledge of homeostasis and disease mechanisms. Array-based spatial transcriptomics (ST) is an unbiased and high-throughput approach to map genes within their spatial context. ST has been applied to chart the organizational landscape of tissues and diseases, such as prostate and pancreatic cancer^1,2^, melanoma^3^, amyotrophic lateral sclerosis^4^ or the developmental human heart^5^. Furthermore, recent studies successfully implemented ST to define the spatial topography of the human dorsolateral prefrontal cortex and its association with schizophrenia and autism^6^.

Several technologies enable the spatial indexing of transcripts and the subsequent mapping of gene expression profiles, their main trade-off being a loss of single-cell resolution. Here, transcripts detected at capture locations (spots) are generally sampled from a mixture of cells which may be homo or heterogeneous. While widely used microarray-based ST techniques utilize 50-100 um spot diameters^7,8^ (10-20 cells, Stahl P et al 2016, 10x Genomics Visium), bead array-based methods further minimized spot sizes to capture cell locations more precisely (2-10 um; Vickovic S et al 2019, Rodrigues SG et al 2019)^9,10^. On the other hand, single-cell RNA sequencing (scRNA-seq) enables the profiling of thousands of single-cell transcriptomes without preserving the spatial context and potentially introducing recovery biases of cell composition. Successful integration of both data modalities could enable an in-depth study of tissue and organ architecture, elucidate cellular cross-talk, spatially track dynamic cell trajectories, and identify disease-specific interaction networks (e.g. between tumors and their microenvironment). Intersecting cell-type-specific genes from scRNA-seq with ST capture sites previously identified local enrichments, sufficient to segment tumor sections into normal and cancerous areas^2^. However, while such analysis allowed predicting the presence or absence of cell types, it lacked the resolution to quantitatively infer cellular compositions at each capture site.

Here we present SPOTlight, a deconvolution algorithm that builds upon a non-negative matrix factorization (NMF) regression algorithm which was previously applied to ST data^10^. Importantly, SPOTlight adds prior information to the model, initializing both the basis and coefficient matrices with cell type marker genes, thereby greatly improving sensitivity and robustness. SPOTlight also relies on non-negative least squares (NNLS) to populate the coefficient matrix of capture locations as well as to determine a spot’s composition. The latter is carried out by defining cell type-specific topic profiles, the distribution of genes defining a cell type or state, and by identifying the weights needed to reconstruct a spot profile. A unit-variance normalization step enables both paired or unmatched ST and scRNA-seq raw count matrices as input. We confirmed the sensitivity and accuracy of SPOTlight predictions on synthetic spots, testing scRNA-seq references of varying qualities (protocols, sequencing depth, cell numbers). SPOTlight showed excellent classification metrics even with low cell and molecule inputs. The possibility to integrate unpaired ST and scRNA-seq data enabled an automated, data-driven interpretation using large reference single-cell atlases, exemplified here using an adult mouse brain atlas^11^. The automated interpretation of ST from patient sections has the potential to digitize pathology and improve patient stratification. As a proof-of-concept, we applied SPOTlight on pancreatic adenocarcinoma (PDAC) data and determined the spatial organization of clinical-relevant immune cell states in the tumor microenvironment.

## Results and Discussion

At the core of SPOTlight, we identify cell type-specific topic profiles used to deconvolute ST spots (**Figure 1**). We set out to use NMF to obtain topic profiles due to its previous success in identifying biologically relevant gene expression programs^12^, as well as its previous implementation in ST analysis^10^. Its non-negative constraint allows it to model count data, which provides more interpretable results than matrix factorization. We seed the model with prior information, guiding it towards biologically relevant results and greatly improving the consistency between runs. Gene expression counts are used as input after a unit variance normalization (by gene) is performed to standardize discretized gene expression levels^10,12^. Importantly, the NMF is initialized by the two main matrices: the basis matrix (W) with unique cell type marker genes and weights, and the coefficient matrix (H) in which each row is initialized, specifying the corresponding relationship of a cell to a topic (i.e. association with a cell type, **Figure 1**). Factorization is then carried out using non-smooth NMF^13,14^. This step returns sparser results during the factorization, promoting cell type-specific topic profiles, while reducing overfitting during training. After factorization, we obtain cell type-specific topic profiles from the coefficient matrix and generate consensus topic signatures across all cells. Subsequently, NNLS regression is used to map each spot’s transcriptome to a topic profile distribution using the unit-variance normalized ST count matrix and the basis matrix previously obtained. Lastly, NNLS is again applied to determine the weights for each cell type that best fit each spot’s topic profile by minimizing the residuals. We use a minimum weight contribution threshold to determine which cell types are contributing to the profile of a given spot, also considering the possibility of partial contributions. NNLS also returns a measure of error along with the predicted cell proportions, allowing the user to estimate the reliability of predicted spot compositions.

**Figure 1.**
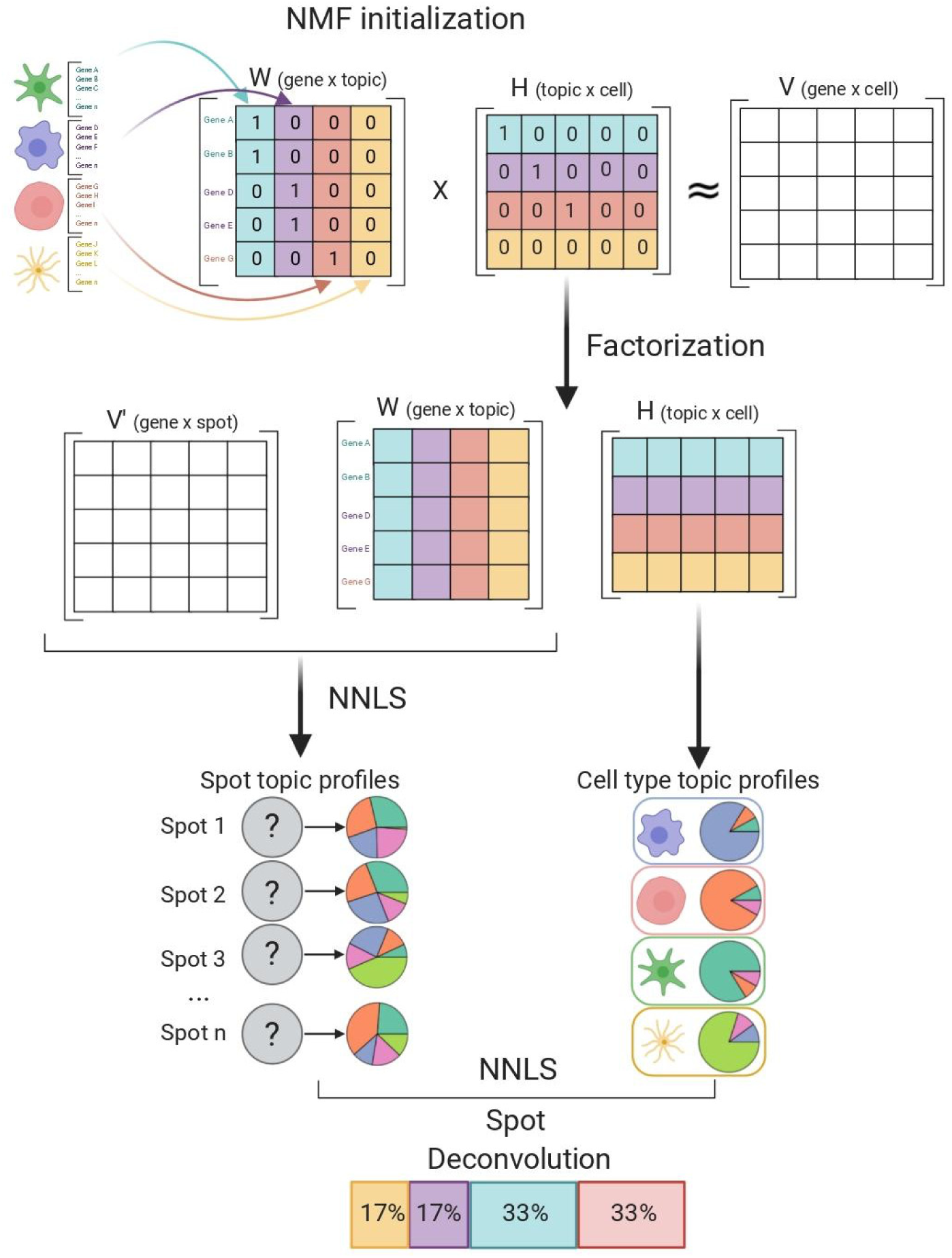
SPOTlight scheme. Step-by-step illustration of SPOTlight’s algorithm. At the beginning of this process we have a count matrix, V, for scRNAseq data and a set of marker genes for the identified cell types. First, we use prior information to initialize the basis and coefficient matrices, W and H respectively. We assume the number of topics, k, to be equal to the number of cell types in the dataset. Each topic is then associated with a cell type; columns in W are initialized with marker genes for the associated cell type with that topic, while rows in H are initialized with the membership of each cell to its associated topic. Second, we proceed with the matrix factorization from which we obtain gene distributions for each topic in W, and topic profiles for each cell in H. Third, we use W to map the ST data, V’, by means of non-negative least squares (NNLS) to obtain H’. Columns in H’ represent the topic profile for each spot. Fourth, from the H matrix obtained from the scRNAseq data we consolidate all the cells from the same cell type to obtain cell type-specific topic profiles. Lastly, we use NNLS to find which combination of cell type-specific topics resembles each spot’s topic profile.

### Benchmarking SPOTlight performance

To evaluate the SPOTlight’s performance, we benchmarked parameters and tested different scenarios with synthetically generated mixtures of cells of known cell type composition. To generate synthetic spots, we selected cells from peripheral blood mononuclear cell (PBMC) scRNA-seq datasets and combined their transcriptomic profiles to different proportions (**Online Methods**). PBMC scRNA-seq data have multiple well-characterized and discrete cell populations, providing an ideal input for benchmarking purposes. Synthetic spots then served as ground-truth to evaluate SPOTlight’s performance to predict cell types and spot composition using the following parameters: *sensitivity* (correctly predicted cell type presence); *specificity* (correctly predicting absence); *precision* (performance when calling a cell type present); *accuracy* (percentage of correctly classified cell types); and *F1 score* (integrating recall/sensitivity and precision). To assess the similarity between the real and predicted proportions, we used the Jensen-Shannon Divergence (JSD), a distance metric that determines the similarity between two probability distributions. As JSD is a distance metric, values closer to 0 signify a higher similarity between both distributions.

When testing the performance on synthetic spots, we obtained a sensitivity of 0.911, an accuracy of 0.78, and an F1 score and specificity of 0.77 and 0.63, respectively. Median JSD values of 0.160 [CI:0.096-0.224] indicated a high accuracy of estimated cell type proportions. The benchmarking results are in line with results from subsequent applications of SPOTlight in different biological scenarios, such as brain tissue or PDAC patient samples; SPOTlight sensitively detected cell types and subtle cell states at their expected locations. A major challenge of NMF is its stochastic nature, requiring repeated iterations from different starting points in order to obtain valid results. To overcome this inherent variability, we initialized the basis and coefficient matrices by seeding them with prior information. Consequently, multiple iterations with seeded NMF regression obtained very similar results for synthetic cell type mixtures (JSD scores, **Figure 2-a-d**). In line with these results, topic profiles from different cell types displayed consistent profiles in all iterations (**Supplementary Fig. 1**) and single cells used to train the model presented comparable topic profiles (**Supplementary Fig. 2**).

**Figure 2.**
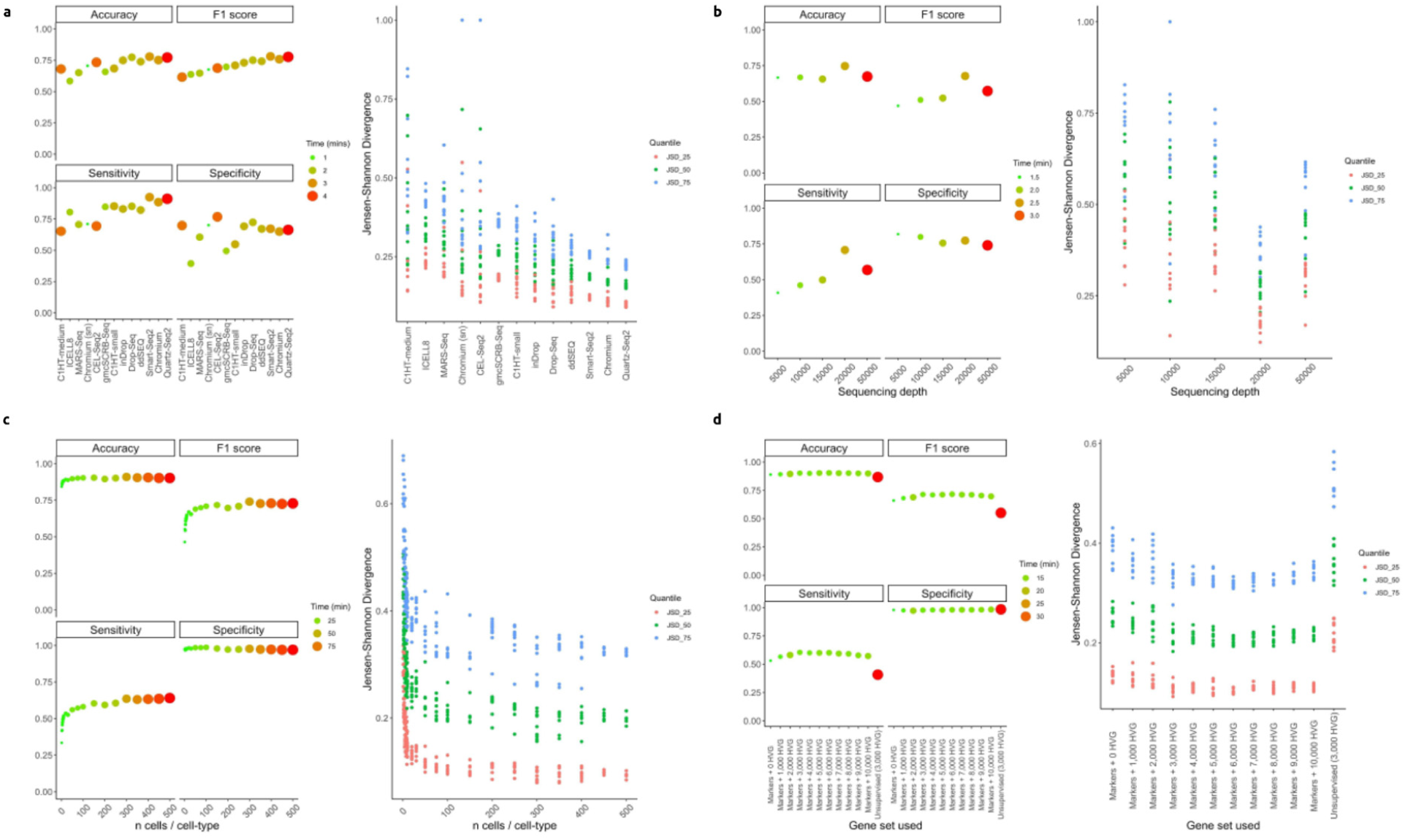
Benchmarking SPOTlight under different technical conditions and parameter optimization. In the classification metrics, the mean of 10 iterations is shown; when assessing Jensen-Shannon Divergence we show the results for each iteration. **a**, Performance of SPOTlight was assessed under different scRNA-seq protocols (Quartz-Seq2, Chromium, and Smart-Seq2) to obtain the best performance. **b**, Benchmarking SPOTlight on the same dataset downsampled to different depths. Performance improved at higher sequencing depths but no sharp decrease at shallow sequencing depths was observed. **c**, Optimizing parameter number of cells per cell type to use to train the model. Peak performance was obtained using 100 cells per cell type, less cells decreased performance while more increased computational time without improving performance. **d**, Optimizing the gene set used to train the model. Optimal performance was obtained when using the union gene set between marker genes and 3,000 highly variable genes (HVG). The unsupervised approach using only the 3,000 HVG performed the worst.

We reason that different input qualities (transcriptome complexity), quantities (cell numbers), and proportion (**Extended Discussion**) would critically impact the performance of SPOTlight. Therefore we opted to test different input scenarios, including scRNA-seq protocols, sequencing depth, cell numbers, and other tunable parameters to simulate variable experimental designs and to identify ideal inputs and limitations of the tools.

We previously benchmarked scRNA-seq protocols for their performance in producing complex sequencing libraries and their suitability to generate reference cell atlases^15^. First, we assessed if the scRNA-seq technologies used to generate data affected the performance of SPOTlight. Different protocols produced vastly variable data qualities and we expected this to impact downstream applications, such as deconvolution algorithms. We used downsampled scRNA-seq datasets (20,000 reads per cell) and trained the SPOTlight model on synthetic mixtures for each protocol (**Figure 2-a**). The best performance was achieved with Quartz-Seq2, Smart-Seq2, and Chromium protocols that also showed excellent benchmarking performance. It is worth highlighting the performance of single-nucleus (sn) sequencing in this context (Chromium sn), which resulted in deconvolution metrics that were comparable to scRNA-seq despite the sampling from a reduced transcriptome pool. In general, scRNA-seq with defined clusters and cell type-specific markers are ideal for optimal performance of SPOTlight. However, other commonly used sc/snRNA-seq protocols also return accurate predictions.

Second, we benchmarked the impact of reduced sequencing depth to identify the performance peak for a cost-effective reference atlas generation. An increased sequencing depth enables the detection of more molecules and genes, including lowly expressed transcripts. When testing SPOTlight on step-wise downsampled datasets (5,000 – 50,000 reads per cell), we observed a critical drop in performance at lower sequencing depth (**Figure 2-b**). While accuracy and specificity were comparable to deeply sequenced datasets, the sensitivity and accuracy of estimated cell type proportions (JSD index) was reduced at lower depths. Nevertheless, despite the lower sensitivity, shallowly-sequenced data such as large atlas projects^16,17^ are also suitable inputs for accurate localization of cell types in space. We detected a peak in performance around 20,000 reads per cell; this sequencing depth was also identified to be most cost-efficient for high-throughput scRNA-seq protocols^18^.

Third, scRNA-seq protocols scale differently with droplet-based methods processing up to millions of cells, while plate-based protocols (e.g. Smart-seq2) generally generate datasets for a few thousand cells. We assessed the impact of input cell numbers on SPOTlight’s performance, also considering computational time as an important factor. We found that the cell number per cell type to train the model was a key parameter (**Figure 2-c**). The optimal value to strike a balance between deconvolution performance and computational time was around 100 cells. Selecting fewer cells would decrease computational time, but the performance has not plateaued. Selecting more cells would drastically increase computational time with marginal improvements on performance. As 100 cells per cell type are in the range of both droplet- and plate-based methods, SPOTlight is suitable for the most commonly used formats of scRNA-seq data.

To train the model, different sources (gene selection) can be used as input. The selection of highly variable genes (HVG) has been shown to be critical for the clustering of scRNA-seq data and we reason that it could also be crucial for spot deconvolution. We further quantified the improvements due to the addition of cell type gene markers, a main difference to previous tools using NMF on ST data^10^. We found that SPOTlight’s performance was optimal when combining both HVG and specific cell types markers to seed the model (**Figure 2-d**). Marker genes critically improved all metrics compared to an unsupervised approach using the 3,000 HVG alone as proposed by the original NMF regression documentation^10^. The number of HVG used had a marginal impact on the performance; however, optimal performance was observed using gene markers combined with the 3,000 HVG.

### Deconvoluting ST derived mouse brain tissue

To validate the SPOTlight performance on complex tissue architectures, we used mouse brain sections, a thoroughly cataloged tissue, presenting well-defined structures, and a plethora of cell types and states with specific molecular fingerprints. As a reference, we used scRNA-seq datasets (Smart-seq2) derived from multiple cortical areas as well as the hippocampus^11^ (∼76.000 cells and 47 annotated cell types/states; **Supplementary Table 1, Supplementary Fig. 3**). To anatomically match the sampling site, we analyzed ST data of the adult mouse brain obtained from anterior and posterior sagittal slices^19^. Two biological replicates for each slice were analyzed to test the robustness of SPOTlight predictions. To validate the predicted spatial cell type distribution within brain areas, we used canonical cell type gene markers along with *in situ hybridization* (ISH) images with cell-level resolution^20^ (Allen Mouse Brain Atlas).

SPOTlight spatial deconvolution of the mouse brain ST data accurately reconstructed the layered and segmented structure of brain anatomy (**Figure 3-a**). The predicted localization of the 47 annotated clusters confirmed their enrichment in distinct layers (e.g. cortical areas) or specific regions (e.g. hippocampus) of the mouse brain (**Supplementary Fig. 4** and **Extended Discussion**). The joint analysis of brain cell types and states resulted in a high-level segmentation, but also provided more detailed information about heterogeneity (composition) of specific areas. A closer inspection confirmed the regional enrichment of specific cell types on their known structures, confirming the high accuracy and sensitivity of the SPOTlight predictions. The results on independent anterior and posterior sections also reflected robust predictions (**Supplementary Fig. 5**).

**Figure 3.**
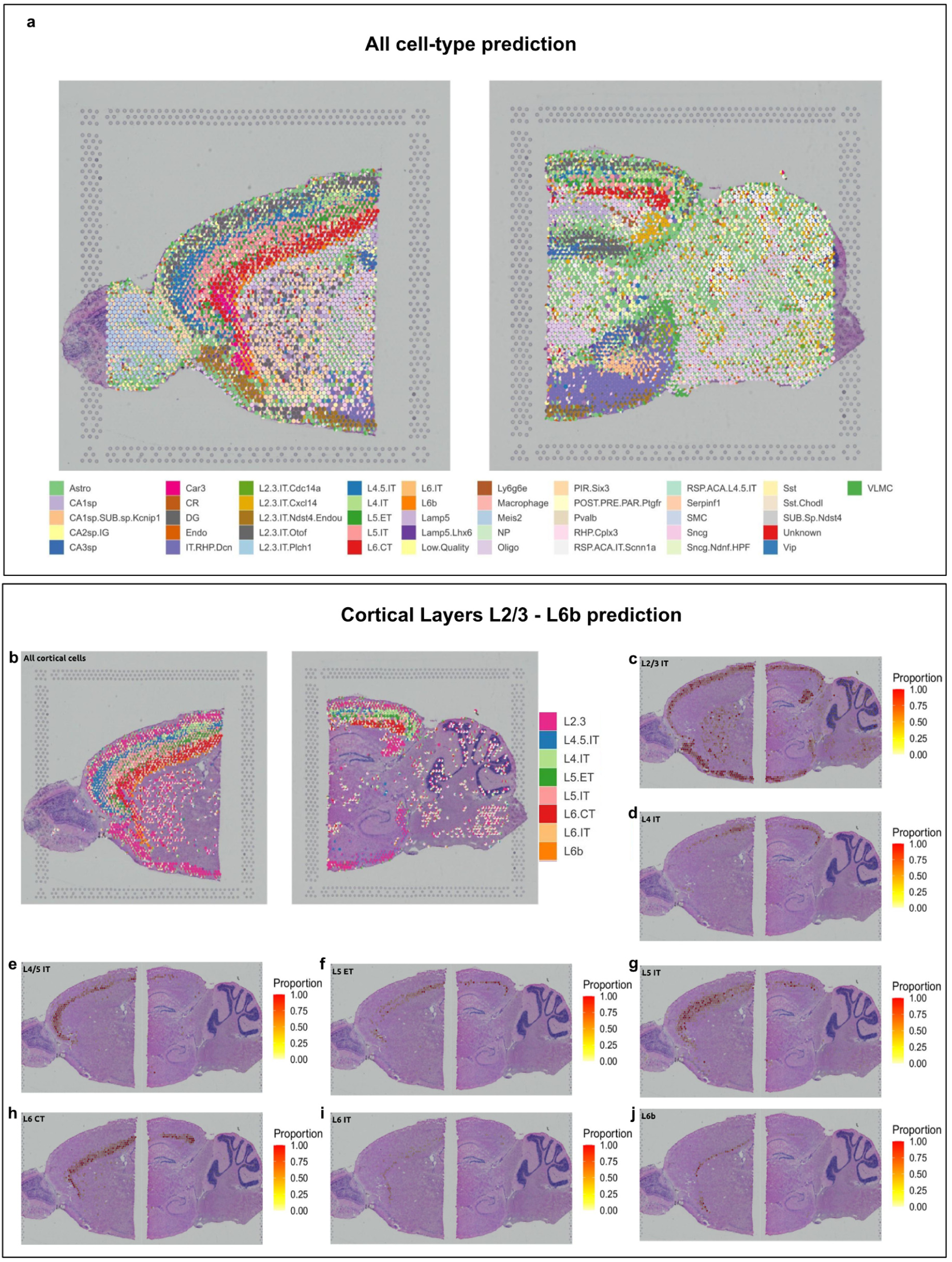
Cell type mapping on sagittal adult mouse brain anterior and posterior slices. **a**. Spatial scatter pie plot representing the proportions of the cells from the reference atlas within capture locations in the adult mouse brain; we can observe the substructures of anatomical regions in the brain as defined by their specific cell types. **b**. Proportions of the cortical cells from the reference atlas within capture locations; SPOTlight is able to capture the cortical structure being able to discern between highly similar neuronal cell types. **c-j**, Proportion within each capture location of each specific cortical neuron type.

Illustrative examples include the SPOTlight deconvolution to delineate the spatial organization of different cortical layers, L2/3 to L6, including layer-specific neuronal subtypes (**Figure 3-b**). Consistent with the strictly layered structure of the cortex, subpopulations aligned along stretched areas descending towards the center (L2-L6, (**Figure 3-c-j**). L6 contributed multiple neuronal subtypes that were all accurately predicted to the respective layer substructure (**Figure 3-h-j**). The ability to differentiate between cortical neuronal subtypes underlines the tool’s sensitivity when similar cell types and states are present in complex tissues.

The hippocampus architecture was first delineated using canonical markers: Cornu Ammonis 1 stratum pyramidale (CA1sp), *Fibcd1*; Cornu Ammonis 2 stratum pyramidale (CA2sp), *Ccdc3*; Cornu Ammonis 3 stratum pyramidale (CA3sp), *Pvrl3*; and Dentate gyrus (DG), *Prox1*^21^. With SPOTlight, we could clearly discern between CA1sp, CA2sp, CA3sp, and the DG (**Supplementary Fig. 6-a,d,g,j**) which was subsequently confirmed by ISH images (**Supplementary Fig. 6-c,f,i,l**). Gene expression measurements of cell type markers from ST alone provided noisy signals (CA1sp, CA2sp, DG) or complete absence (CA3sp) (**Supplementary Fig. 6-b,e,h,k**) related to the sparsity of ST data; highlighting the need for more sophisticated spatial annotation tools.

### Charting spatial heterogeneity in human cancer

To further validate a broader application spectrum and to test its performance in complex human tissues, we applied SPOTlight on ST data from PDAC patient samples^2^, generated with a different ST protocol version than the mouse brain data^9^ (**Extended Discussion**). Sample-matched scRNA-seq data (inDrop) was analyzed to chart the tumor composition and subsequently used to train the SPOTlight model (**Figure 4-a**). When integrating scRNA-seq and ST (PDAC-A), we observed a discrete regional enrichment of normal pancreatic and neoplastic cell types (**Figure 4-b**). In detail, normal cell types of the pancreas were mainly excluded from the tumor fraction and further split into acinar and ductal areas. Centroacinar ductal populations appeared in the duct epithelium, while terminal ductal populations were found in both duct epithelium as well as co-localizing in the cancerous part of the tissue (**Supplementary Fig. 7**). In line with previous results^2^, we detected the intermixing of two distinct tumor cell clones and the enrichment of a ductal population with a hypoxia gene signature in the cancerous region (**Figure 4-c**).

**Figure 4.**
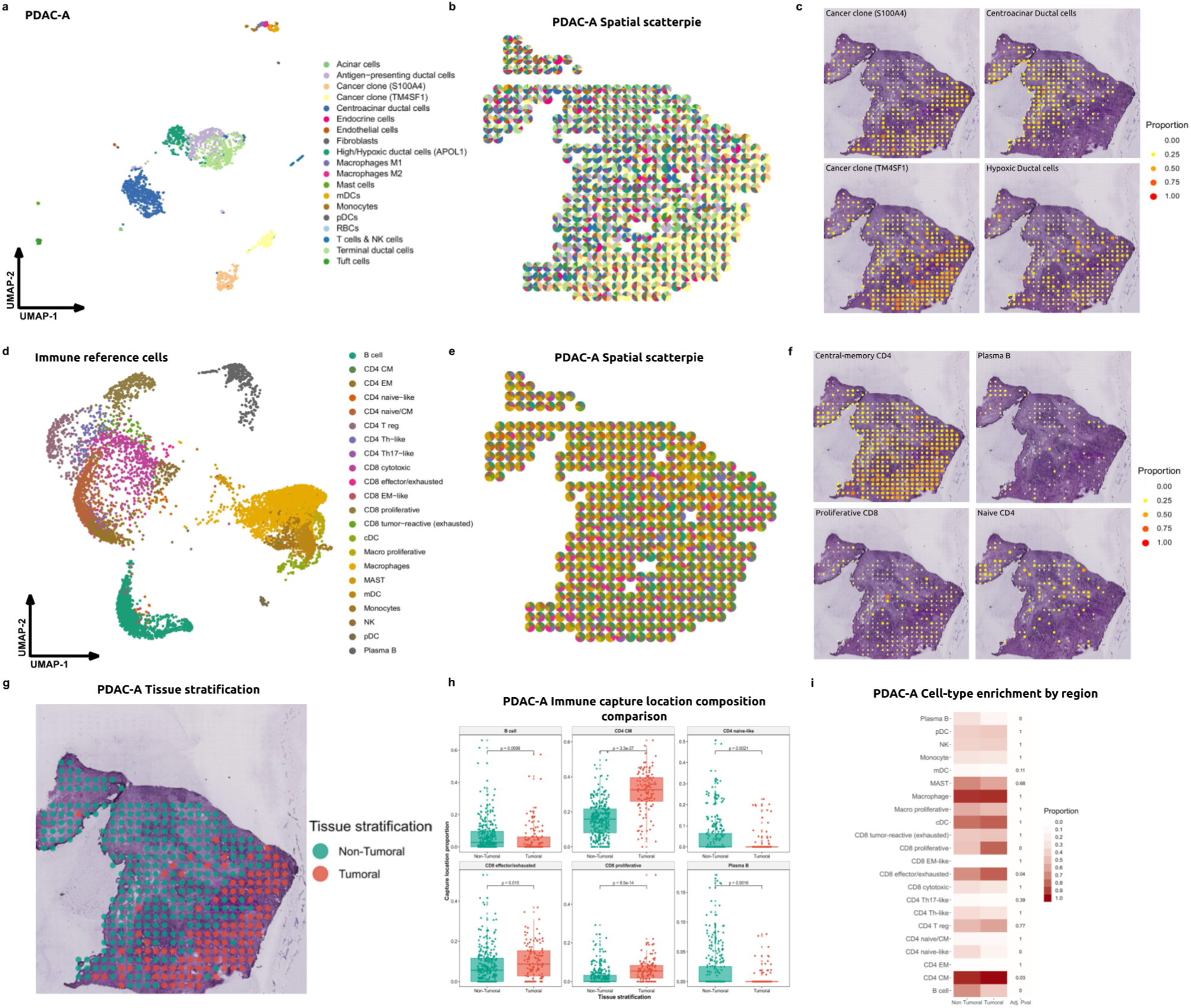
Mapping cell subpopulations across the tissue, charting tumoral and immune cell distribution on the tissue to identify differential immune microenvironments in tumoral vs non-tumoral regions. **a**, UMAP projections of 1,926 cells from PDAC-A, paired data from tissue slices. Cells are colored and labelled according to the cell type annotations from the original paper. **b**, Spatial scatter pie plot representing the proportions of the cell types in the paired inDrop dataset within the capture locations. **c**, Predicted proportion within each capture location for cancer clones S100A4 and TM4SF1 and centroacinar and hypoxic ductal cells. **d**, UMAP projections of pancreatic immune reference cells mapped onto PDAC-A ST1. **e**, Spatial scatter pie plot representing the proportions of the immune cells within the capture locations. **f**, Predicted proportion within each capture spot for central memory and naive CD4, Plasma B cells and proliferative CD8 cells. **g**, Tissues tratification by tumoral – non-tumoral capture locations, stratification coincides with pathologist’s annotation. **h**, Cell type proportion comparison within each spot between tumoral and non-tumoral sections. Central memory CD4, proliferative and effector CD8 cells are found at higher proportions within the tumoral region **i**, Proportion of capture locations containing each immune cell type within the tumoral and non-tumoral sections.

To shed light on the distribution of immune cells in the tumor sections, we integrated, clustered, and annotated an external single-cell PDAC dataset with a specific focus on the tumor immune microenvironment^22^. Briefly, scRNA-seq data from 24 PDAC patients and 41,986 cells were merged to identify a total of 10,623 immune cells (**Supplementary Table 2**). Clustering and curated annotation resulted in 22 immune subpopulations with 12 T-cell, 3 macrophage/monocyte, 2 B-cell, 4 dendritic, and 1 MAST cell clusters (**Figure 4-d**). SPOTlight trained on PDAC immune cells and applied on the PDAC-A ST slides resulted in a remarkable local enrichment of tumor-specific cell states (**Figure 4-e,f**, and **Supplementary Fig. 8**). In line with the regional distribution of normal and cancer cells, we identified a striking segmentation of immune cell states in the PDAC section (**Figure 4-g** and **Supplementary Fig. 9**). While naive CD4 and plasma B-cells were enriched in the normal pancreas tissue, central-memory CD4 T-cells as well as effector/exhausted and proliferative CD8 cells were significantly increased in the tumor (p < 0.01, **Figure 4-h,i** and **Supplementary Fig. 9**). In a second PDAC patient section (PDAC-B), central-memory CD4 and effector CD8 cells again co-localized with the tumor areas, while plasma B-cells were depleted from that area and mainly found together with endothelial and endocrine cells (**Supplementary Fig. 10**). Most importantly, the enrichment of central-memory CD4 cells could not be detected through their presence alone. While the PDAC-B case showed an exclusive localization to the tumor area, central-memory CD4 cells were highly abundant in all areas, but to higher proportions in the tumor in PDAC-A. This finding strongly underlines the need to sensitively deconvolute spot composition to enable precise pathology assessments. The regional differences and local immune cell enrichments further allowed us to compute cell-cell interaction networks using the cell’s co-localization in the PDAC sections (**Supplementary Fig. 11)**. Such visualization underlined the concerted interaction of tumor-resident immune cells and could provide further insight into the peculiarities of tumor microenvironments.

## Conclusion

SPOTlight proved to be a robust, accurate, and sensitive tool to determine cell-type locations and a fine-grained composition of ST spots. We showed that scRNA-seq quality can impact its performance, obtaining the best results with deeply sequenced data from complex sequencing libraries. Nevertheless, SPOTlight also returns accurate predictions with shallowly sequenced references; an important feature when using large atlas projects as a reference. We further showed that as few as 100 cells per cell-type were sufficient to train the model without prolonged computation time. Applying SPOTlight on vastly different biological scenarios, different technology versions, and using matched and external references confirmed its broad and flexible application spectrum. This makes it a universal tool to combine both pillars of the single-cell genomics field^23^ and to deduce cellular function and organization *in situ*. We are particularly excited about the potentially transformative impact on pathological assessments. Using an external immune reference to delineate the localization of immune cells in tumors could be implemented in automated digital pathology systems, where query ST patient samples are screened for immune cell composition and distribution. Importantly, both features have been related to patient prognosis and (immuno-) therapy response. Thus, we foresee spatial deconvolution using SPOTlight or similar tools to have a major impact on future cancer patient management and on precision oncology.

## Acknowledgment

This work has received funding from the Ministerio de Ciencia, Innovación y Universidades (SAF2017-89109-P; AEI/FEDER, UE). This publication is part of a project (BCLLATLAS) that has received funding from the European Research Council (ERC) under the European Union’s Horizon 2020 research and innovation programme (grant agreement No 810287). This project has been made possible in part by a grant from the Chan Zuckerberg Initiative. We acknowledge support of the Spanish Ministry of Science and Innovation to the EMBL partnership, the Centro de Excelencia Severo Ochoa and the CERCA Programme / Generalitat de Catalunya. We also acknowledge the support of the Spanish Ministry of Science and Innovation through the Instituto de Salud Carlos III, the Generalitat de Catalunya through Departament de Salut and Departament d’Empresa i Coneixement, and the Co-financing by the Spanish Ministry of Science and Innovation with funds from the European Regional Development Fund (ERDF) corresponding to the 2014-2020 Smart Growth Operating Program.

## Availability of data and code

The SPOTlight code and the analysis notebooks to reproduce the aforementioned analysis are hosted at https://github.com/MarcElosua/SPOTlight and https://github.com/MarcElosua/SPOTlight_deconvolution_analysis.

The ST and scRNA-seq data has been previously published^11,15,22^ and is freely available at the Gene Expression Omnibus (GEO) under GSE133549, and GSE71585 and GSE111672 and n the Genome Sequence Archive under project PRJCA001063.

Docker environments are available for R and Rstudio at Docker Hub marcelosua/spotlight_env_r:latest and marcelosua/spotlight_env_rstudio:latest respectively.

## Author’s Contributions

HH and ME designed the study. ME developed SPOTlight and performed ST and scRNA-seq data analyses. PN compiled the PDAC scRNA-seq data and performed clustering and immune cell annotation. EM and IG supported the data analysis. HH and ME wrote the manuscript. All authors read and approved the final version.

## Extended Discussion

### Benchmarking SPOTlight performance

The minimum weight contribution a cell type required to be considered as present within a SPOT was acceptable between 0.06-0.09 and showed optimal performance at 0.07 (F1 score = 0.798; **Supplementary Fig. 12**). Higher and lower thresholds showed a clear trade-off between specificity and sensitivity. In practical terms, this means that cell types are included, which contribute at least 6-9% of the spots’ topic profiles. This seems ideal considering that widely used microarray-based ST spots contain around 10 cells. Thus, lowly abundant subpopulations with only single cells contributing to a spot’s profile can be readily identified by SPOTlight.

### Deconvoluting ST derived mouse brain tissue with SPOTlight

In mouse brain tissue slices, we identified spatial locations for different cell types from which prior knowledge of their spatial topography is available. When focusing on the hippocampus, CA1sp, CA2sp, CA3sp and DG were predicted to be located on their respective structures^21^ (**Supplementary Fig. 6**). Being able to differentiate similar cell types showed the robustness and sensitivity of the model and suggests its broad utility in other complex tissues. However, the model also predicted L2/3 and CA1sp neurons to be spatially located in the interior part of the brain as well as in the cerebellum. Of note, the reference scRNA-seq data used to train the model were sampled from cortical and hippocampal regions only. No specific cell populations profiles from the other regions were available. Thus, neurons with similar transcriptomic profiles not present in the training set could explain the signal observed within these regions. Therefore, it is important to note that in order to accurately map the cell type composition a representative single-cell sample is required.

It is of note that three of the clusters in the scRNA-seq training data were labelled as low-quality, unknown, and doublets (mixture of multiple transcriptomes). When predicting their spatial location, we observe that doublets were not predicted in the ST slides and low-quality/unassigned cells displayed only low, residual and scattered presence (**Supplementary Fig. 4,5**). These results give strength to the robustness and specificity of the model with technical artifacts not showing local enrichments or specific structures. It is of note though that the low-quality cluster had a topic profile similar to Vip cells, suggesting an enrichment of low-quality cells from this cell type (**Supplementary Fig. 1**).

### Charting spatial heterogeneity in human cancer

Testing SPOTlight on PDAC sections also allowed us to test the tool on a different ST technology version, with larger capture locations (100um vs 55um) and a wider center-to-center distance (200um vs 100um). Larger spots capture more cells (between 20 and 100)^2^, allowing to assess SPOTlight’s performance with different prior assumptions. To account for larger capture locations, we kept all optimal parameters, but adjusted the minimum contribution threshold to 0.01 so that cell types contributing ≥1% were detected.

## Online Methods

### Implementation

#### Non-Negative Matrix Factorization Regression

The following annotations will be used when describing the model:

- N – Set of all cells from scRNAseq.
- M – Set of all capture locations from spatial data.
- G – Set of selected genes from scRNAseq, cell type marker genes + 3000 highly variable genes.
- G’ – Set of all genes from spatial data.
- Gi – G∩G’, intersection between G and G’.
- C – Number of cell types in the scRNAseq dataset
- K – Number of topics to use to reduce the dimensionalities, equal to C.
- V – matrix of dimensions Gi x N containing data from scRNAseq
- W – matrix of dimension Gi x K containing the gene distribution for each topic, basis between V and H.
- H – matrix of dimensions K x N containing the topic distribution for each cell.
- V’ – matrix of dimensions Gi x M containing spatial data.
- H’ – matrix of dimensions K x M containing the topic distributions for each capture location.
- Q – matrix of dimension K x C containing the topic distributions for each cell type.
- P – matrix of dimension C x M containing the cell type weights for each capture location.

At the core of our tool, we use Non-Negative Matrix Factorization (NMF) along with Non-Negative Least Squares (NNLS). NMF is used to factorize a matrix into two or more lower dimensionality matrices without negative elements. We first have an initial matrix V, which is factored into W and H. Unit variance normalization by gene is performed in V and V’ in order to standardize discretized gene expression levels, “*counts-umi”*^10,12^. Factorization is then carried out using the non-smooth NMF method^13^, implemented in the R package *NMF*^14^. This method is intended to return sparser results during the factorization in W and H, thus promoting cell-type-specific topic profile and reducing overfitting during training. Before running factorization, we initialize each topic, column, of W with the unique marker genes for each cell type with weights 1 – P value. The marker genes are obtained from Seurat’s function *FindAllMarkers*. In turn, each topic of H is initialized with the corresponding belongance of each cell for each topic, 1 or 0. This way, we seed the model with prior information, thus guiding it towards a biologically relevant result. This initialization also aims at reducing variability between runs and improving the consistency between runs.

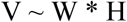

Second, non-negative least squares (NNLS) regression is used to map each capture location’s transcriptome in V’ to H’ using W as the basis. We obtain a topic profile distribution over each capture location which we can use to determine its composition.

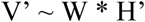

Third, we obtain Q, cell-type specific topic profiles, from H. We select all cells from the same cell type and compute the median of each topic for a consensus cell-type-specific topic signature. We then use NNLS to find the weights of each cell type that best fit H’ minimizing the residuals.

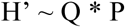

We use a minimum weight contribution to determine which cell types belong within a capture location. 0.09% is set by default, related to the expected number of cells at the capture locations (1-10 cells). In a scenario with 10 cells, we would detect all and also account for partially contributing cells.

By using NNLS, we are able to return a measure of error along with the predicted cell proportions. To do so, we calculate the total sum of squares (TSS) and the residual sum of squares (RSS) for each row. By dividing the RSS by the TSS we obtain the percentage of unexplained residuals for each spot. This measure can be used to assess the quality of a predicted composition.

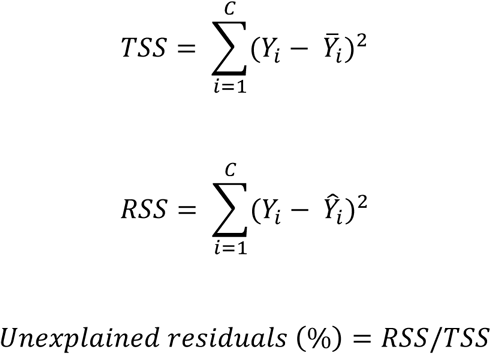

### Parameters

Three important parameters can be adjusted and tuned in order to optimize the performance of this tool: 1) number of cells per cell-type, 2) the supervised vs unsupervised approach along with marker gene sets, and 3) the minimum weight contribution threshold to include a cell as present. We benchmarked these parameters to assess their impact on the performance as follows:

- The number of cells per cell-type used to train the model. We identify the optimal number of cells maximizing performance along with computing time.
- The supervised vs unsupervised approach. For the former, we also tested marker genes together with different numbers of HVG. For the unsupervised approach, we used the 3,000 HVG as in the original NMFreg^10^.
- The minimum weight contribution. This refers to the NNLS weights from C that best fit H’. This weight contribution must be set. Due to the nature of NNLS, there may be cell types contributing a low amount just to residually minimize the squares, and therefore, adding noise to the prediction.

### Synthetic spots

To be able to test the tool’s performance, to benchmark parameters and to apply it on different data types, we generated synthetic mixtures of cells with defined composition. To generate these synthetic test mixtures, we selected between 2 and 8 cells from the scRNA-seq datasets and combined their transcriptomic profiles. If the resulting mixture had >25,000 UMI counts we randomly downsampled it to 20,000 UMI counts in order to better simulate biological capture locations. Test mixtures can be generated using the SPOTlight function *test_spot_fun*.

### Performance evaluation

To address how well the model performed, we assessed several parameters using synthetic and real datasets. From the predicted composition, we first evaluated if we were able to accurately predict when a cell type was correctly predicted within the mixture. Moreover, we also assessed if the predicted proportions were an accurate representation of the true composition. The former is a classification problem for which we used the following parameters; *sensitivity*, if a cell type correctly predicted to be present within the capture location; specificity, predicting its absence when its not present; *precision*, how good we are at identifying cell-types present; *accuracy*, percentage of correctly classified cell types; and *F1 score*, integrating sensitivity and precision. For the latter we used the *Jensen-Shannon Divergence* (JSD) distance metric used to measure the similarity between two probability distributions, P and Q.

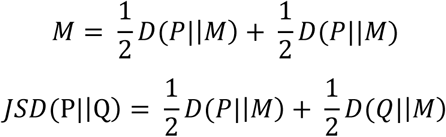

### Benchmarking

For technology benchmarking, we assessed if the technology used to obtain scRNAseq data affected the performance of the model. We used data from peripheral blood mononuclear cells (PBMC) downsampled to the same sequencing depth (20,000 reads/cell). Specifications on how the data was generated and processed can be found elsewhere^15^. For each technology, we trained the model and tested synthetic mixtures of 2-8 cells. We assessed: Cel-Seq2, ChromiumV2, Chromium V2 single-nucleus, C1HT-medium, C1HT-Small, ddSeq, Drop-Seq, gmcSCRB-Seq, ICELL8, inDrop, MARS-Seq, QUARTZ-Seq2, and SMART-Seq2. Data is publicly available through the Gene Expression Omnibus (GSE133549). To benchmark the effect of sequencing depth, we analyzed a Chromium V3 PBMC dataset^15^ (GSE133549) downsampled to different depths using zUMI^24^ (5,000, 10,000, 15,000, 20,000, 50,000 reads/cell). Shared cell types between the different datasets were used excluding biases introduced by varying cell type numbers.

### Mouse brain deconvolution

To assess the tool’s performance on a biological dataset, we used mouse brain as model tissue. Despite its complexity with multiple cell types and states, it presents well-defined structures with location-specific types. We used a mouse brain reference scRNA-seq dataset comprised of cells sampled from multiple cortical areas and the hippocampus, provided by the Allen Institute, with ∼76.000 cells and 47 annotated clusters sequenced using SMART-Seq2^11,25^ (GSE71585). The spatial transcriptomics data of an adult mouse brain (anterior and posterior sagittal slices) was obtained from 10X Genomics^19^. Two replicates for each slice were available and used to confirm the predictions. To validate the predicted cell type spatial distribution within the brain structure, we used known cell-type gene markers along with reference *in situ hybridization* (ISH) images data at cellular-level resolution from the Allen Mouse Brain Atlas^20^. The marker genes used for the hippocampal cell types represented in this study were: Cornu Ammonis 1 stratum pyramidale (CA1sp), *Fibcd1*; Cornu Ammonis 2 stratum pyramidale (CA2sp), *Ccdc3*; Cornu Ammonis 3 stratum pyramidale (CA3sp), *Pvrl3*; and Dentate gyrus (DG), *Prox1*, as reported in Cembrowski M, et al 2016.

In situ hybridization images were obtained from the Allen Brain Atlas. Links to the images are the following:

- *Fibcd1*:mouse.brain-map.org/experiment/siv?id=69672462&imageId=69647545&initImage=ish&coordSystem=pixel &x=4464.5&y=3184.5&z=1
- *Ccdc3*:mouse.brain-map.org/experiment/siv?id=68844056&imageId=68705406&initImage=ish&coordSystem=pixel &x=4352.5&y=2880.5&z=1
- *Pvrl3*:mouse.brain-map.org/experiment/siv?id=69816733&imageId=69747543&initImage=ish&coordSystem=pixel &x=5744.5&y=3576.5&z=1
- *Prox1*:mouse.brain-map.org/experiment/siv?id=69289763&imageId=69177644&initImage=ish&coordSystem=pixel &x=5416.5&y=3720.5&z=1

### Pancreatic ductal adenocarcinoma

We used pancreatic ductal adenocarcinoma (PDAC) ST data publicly available through the Gene Expression Omnibus (GSE111672)^2^. Spatial data for this study was generated with the original spatial transcriptomics technology^9^, while scRNAseq data was generated using inDrops. Further specifications on how the data was generated and processed can be found elsewhere^2^. In total, 10 spatial slides from 6 tumor samples are available, 2 of which (PDAC-A and PDAC-B) have 3 biological replicates and paired scRNAseq data. For the purpose of this study we used samples PDAC-A and PDAC-B and selected sections that harbored both normal and tumor areas (identified through the mapping of normal cell types and tumor clones). The ST data is accessible through GSM3036911 for PDAC-A and, GSM4100723 for PDAC-B, scRNAseq for PDAC-A data through GSM3036909, GSM3036910, GSM3405527, GSM3405528, and PDAC-B data through GSM3405531, GSM3405532, GSM3405533. Filtering and data processing was carried out as specified in the original publication, keeping cells with ≥1000 UMIs, ≤20% mitochondrial transcripts, and ≤30% ribosomal transcripts^2^. In PDAC-B, one cluster of ductal cells with low UMIs and high mitochondrial content was removed.

To generate a comprehensive immune cell type reference atlas for PDAC, we re-analyzed scRNA-seq data from Peng J et al 2019; Genome Sequence Archive: ID PRJCA001063). From this dataset only the tumoral pancreas samples were included. Cells with >20% of mitochondrial content and <100 UMIs were removed. We normalized, scaled, extracted the highly variable genes and performed PCA analysis on the remaining cells prior to clustering. Resulting clusters were annotated according to gene markers provided in the original manuscript. All the tumor and non-immune cells were identified and removed by marker gene analysis. For the detailed annotation of the immune cells, we first used canonical markers to group them into the major cell types (i.e. *CD79A, CD68* and *CD3E* for B-cells, myeloid cells, and T-cells, respectively). To further stratify the cells into cell states, we reclustered and annotated each of them comparing the cluster markers to well-characterized single-cell gene sets of the tumor microenvironment^26–28^ by computing the Jaccard similarity index using *matchSCore2*^15^. We were able to identify all of the expected cell populations, including rare immune cell states.

When stratifying the tissue into tumoral and non-tumoral sections, tumoral spots contained >40% cancer-cell proportion. Cell type proportions within the spots were compared between regions and significance assessed using a non-parametric test (Mann-Whitney). To assess cell type enrichment between regions, we computed the proportion of spots containing each cell type. The significance between the proportions was assessed with a permutations test where the cell type specific statistic distribution was created randomly 10,000 times for each cell type. Moreover, we also assessed a third region, intermediate, between the tumoral and non-tumoral regions. Here, regions were defined as follows: tumoral, >40% cancer-cell proportion; intermediate, <40% cancer-cell and ductal-cell proportion; and non-tumoral, >40% ductal-cell type proportion. Again, cell type proportions within the spots were compared between regions and significance assessed with a Mann-Whitney test. Bonferroni adjusted p-values are reported for multiple comparisons.

### Code versions and availability

This tool is developed to run with R versions ≥3.5; docker images with the appropriate environment are available at Docker hub: marcelosua/spotlight_env_rstudio and marcelosua/spotlight_env_r.

