## Supplementary Figures and Tables. for "SPOTlight: Seeded NMF regression to Deconvolute Spatial Transcriptomics Spots with Single-Cell Transcriptomes"

**Supplementary Table 1: Composition of the Allen Mouse Brain Atlas dataset.**

| Cell type | Number of cells | Cell type | Number of cells | Cell type | Number of cells |
|---|---|---|---|---|---|
| **Astro** | 971 | **L4 IT** | 5455 | **POST-PRE-PAR Ptgfr** | 50 |
| **CA1sp** | 1506 | **L4/5 IT** | 6280 | **Pvalb** | 4071 |
| **CA1sp/SUB-sp Kcnip1** | 189 | **L5 ET** | 1994 | **RHP Cplx3** | 152 |
| **CA2sp/IG** | 20 | **L5 IT** | 3306 | **RSP_ACA IT Scnn1a** | 202 |
| **CA3sp** | 321 | **L6 CT** | 6826 | **RSP/ACA L4/5 IT** | 282 |
| **Car3** | 1997 | **L6 IT** | 4645 | **Serpinf1** | 279 |
| **CR** | 39 | **L6b** | 2538 | **SMC** | 174 |
| **DG** | 2491 | **Lamp5** | 3626 | **Sncg** | 1079 |
| **Doublet** | 813 | **Lamp5 Lhx6** | 966 | **Sncg/Ndnf HPF** | 367 |
| **Endo** | 320 | **Low Quality** | 457 | **Sst** | 5577 |
| **IT RHP Dcn** | 200 | **Ly6g6e** | 253 | **Sst Chodl** | 276 |
| **L2/3 IT Cdc14a** | 298 | **Macrophage** | 185 | **SUB-Sp Ndst4** | 474 |
| **L2/3 IT Cxcl14** | 1241 | **Meis2** | 172 | **Unknown** | 70 |
| **L2/3 IT Ndst4 Endou** | 217 | **NP** | 2271 | **Vip** | 6368 |
| **L2/3 IT Otof** | 6066 | **Oligo** | 237 | **VLMC** | 187 |
| **L2/3 IT Plch1** | 777 | **PIR Six3** | 22 | | |

**Supplementary Table 2: Composition of the PDAC immune reference scRNAseq dataset.**

| Cell type | Number of cells | Cell type | Number of cells | Cell type | Number of cells |
|---|---|---|---|---|---|
| **B cell** | 2429 | **CD8 cytotoxic** | 478 | **MAST** | 131 |
| **CD4 CM** | 3 | **CD8 effector/exhausted** | 26 | **mDC** | 792 |
| **CD4 EM** | 579 | **CD8 EM-like** | 64 | **Monocytes** | 357 |
| **CD4 naive-like** | 48 | **CD8 proliferative** | 393 | **NK** | 96 |
| **CD4 naive/CM** | 679 | **CD8 tumor-reactive (exhausted)** | 68 | **pDC** | 28 |
| **CD4 T reg** | 270 | **cDC** | 153 | **Plasma B** | 412 |
| **CD4 Th-like** | 128 | **Macro proliferative** | 278 | | |
| **CD4 Th17-like** | 65 | **Macrophages** | 3146 | | |

**CM: Central Memory; EM: Effector memory; T reg: regulatory T cells; Th: T helper; cDC: common dendritic cells; mDC: myeloid denritic cells; NK: natural killer; pDC: plasmacytoid dendritic cells.**

**Supplementary Table 3: Composition of the PDAC-A scRNAseq dataset.**

| Cell type | Number of cells | Cell type | Number of cells |
|---|---|---|---|
| **Acinar cells** | 13 | **Macrophages M2** | 21 |
| **Antigen-presenting ductal cells** | 287 | **Mast cells** | 14 |
| **Cancer clone (S100A4)** | 170 | **mDCs** | 45 |
| **Cancer clone (TM4SF1)** | 126 | **Monocytes** | 18 |
| **Centroacinar ductal cells** | 529 | **pDCs** | 13 |
| **Endocrine cells** | 3 | **RBCs** | 15 |
| **Endothelial cells** | 11 | **T cells & NK cells** | 40 |
| **Fibroblasts** | 5 | **Terminal ductal cells** | 350 |
| **High/Hypoxic ductal cells (APOL1)** | 215 | **Tuft cells** | 32 |
| **Macrophages M1** | 19 | | |

**mDCs: myeloid denritic cells; pDCs: plasmacytoid dendritic cells; RBCs; red blood cells**

**Supplementary Table 4: Composition of the PDAC-B scRNAseq dataset.**

| Cell type | Number of cells | Cell type | Number of cells |
|---|---|---|---|
| **Acinar cells** | 6 | **Mast cells** | 13 |
| **Antigen-presenting ductal cells** | 211 | **mDCs** | 35 |
| **Cancer clone (TM4SF1)** | 339 | **Monocytes** | 20 |
| **Centroacinar ductal cells** | 152 | **RBCs** | 3 |
| **Endocrine cells** | 13 | **Terminal ductal cells** | 736 |
| **Endothelial cells** | 159 | **Tuft cells** | 37 |
| **Macrophages** | 9 | | |

**mDCs: myeloid denritic cells; RBCs; red blood cells**

Supplementary Fig. 1

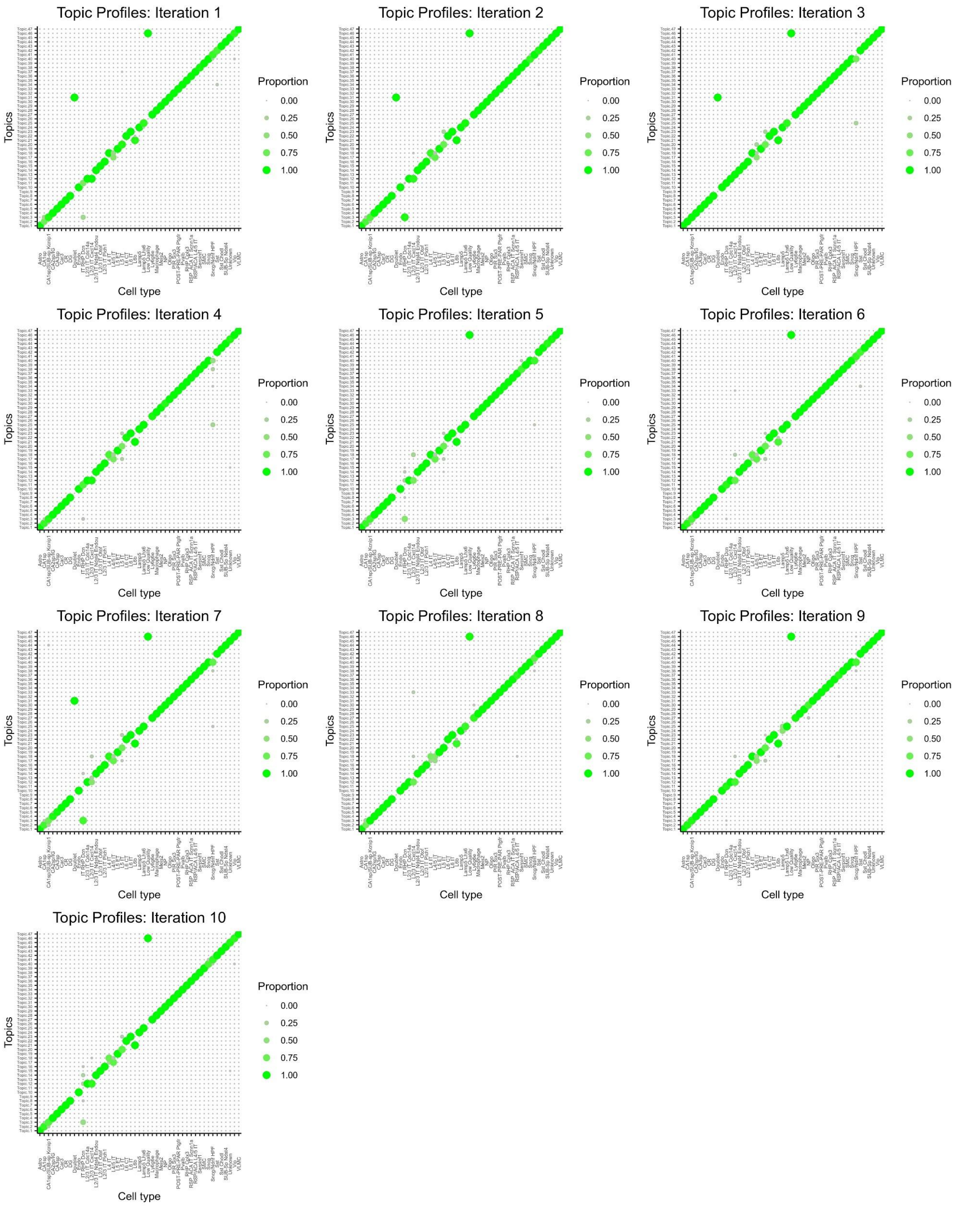


**Supplementary Figure 1 |** Cell type specific topic profiles obtained from 10 random iterations, cell-type topic profiles are consistent across multiple iterations.

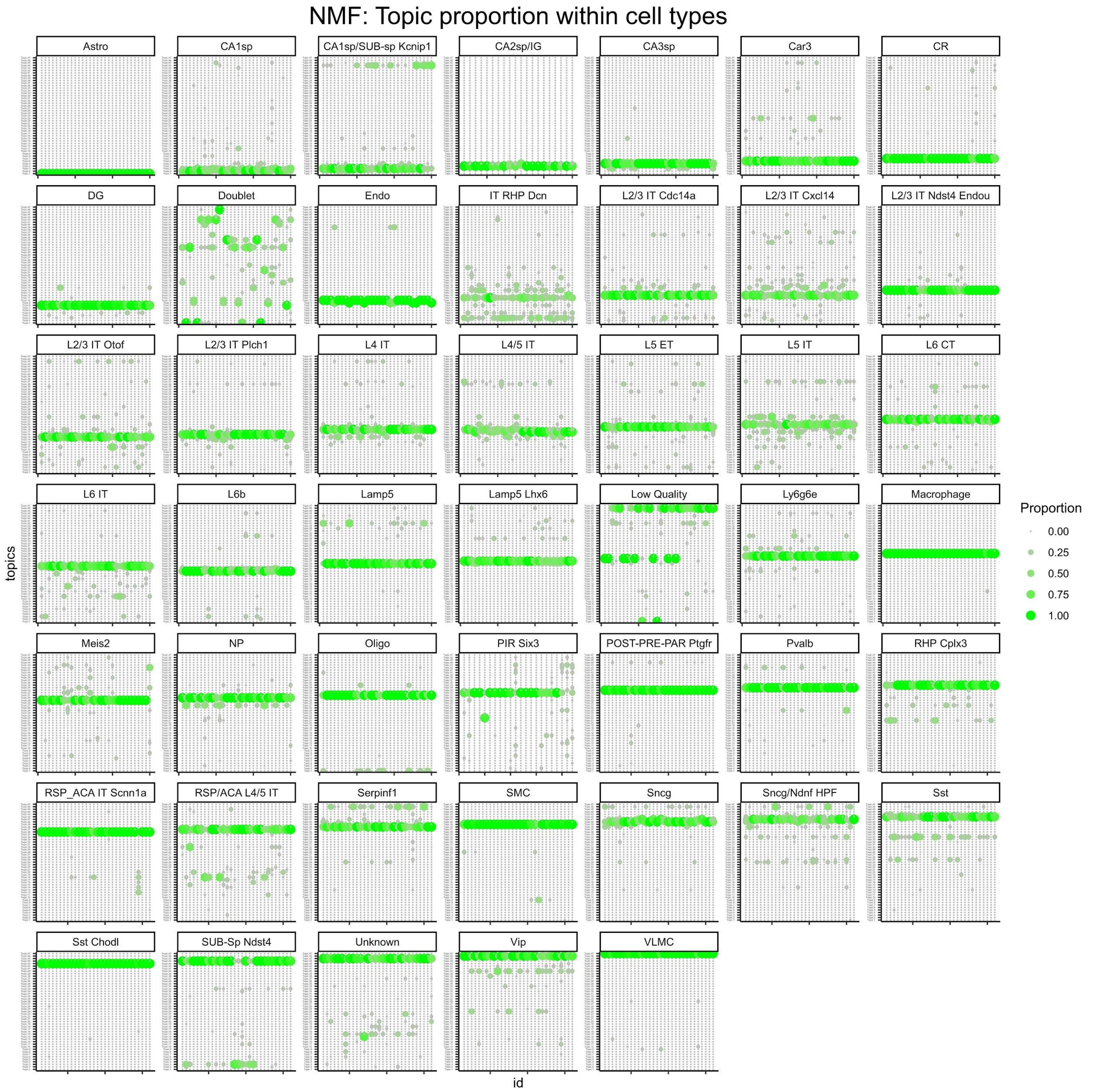


**Supplementary Figure 2 |** Topic profiles of individuals cells used to train the model by cell type, cells from the same cell type had very similar profiles.

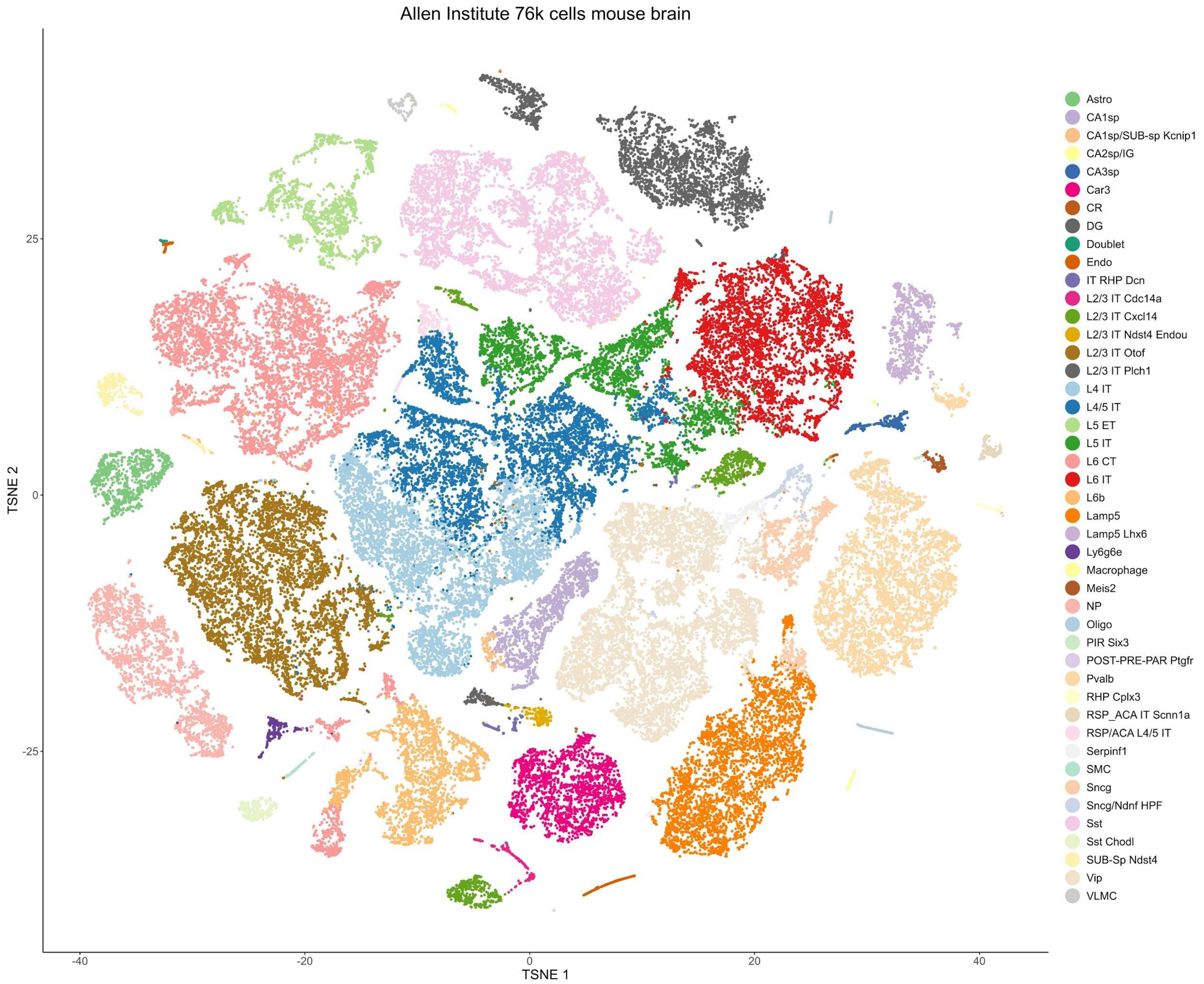


**Supplementary Figure 3 |** T-SNE projections of 74.967 cells from the Allen Institute mouse brain scRNAseq reference atlas. Cells are colored and labelled according to the cell type annotations provided.

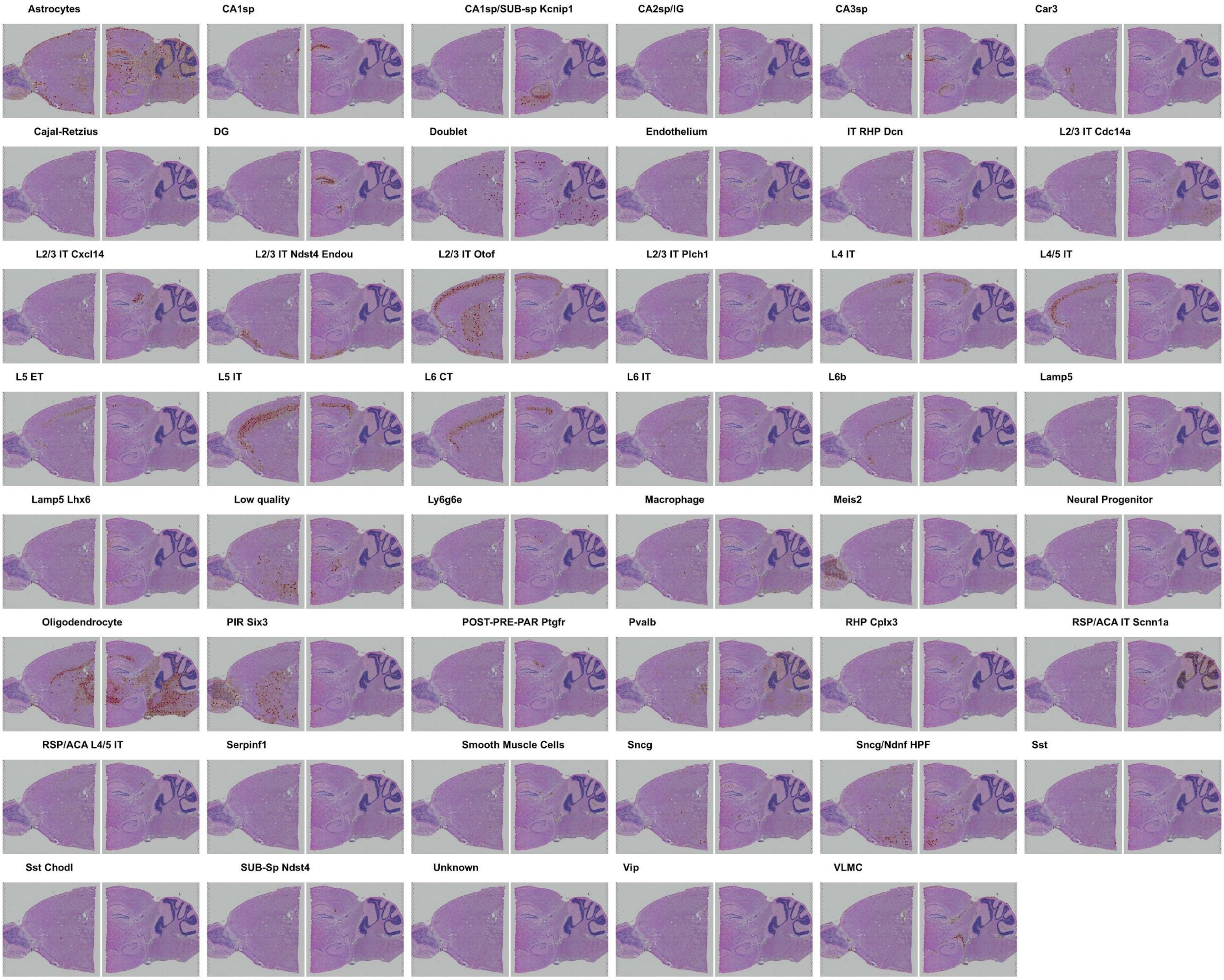


**Supplementary Figure 4 |** Predicted cell-type proportion within each capture location replicate 1.

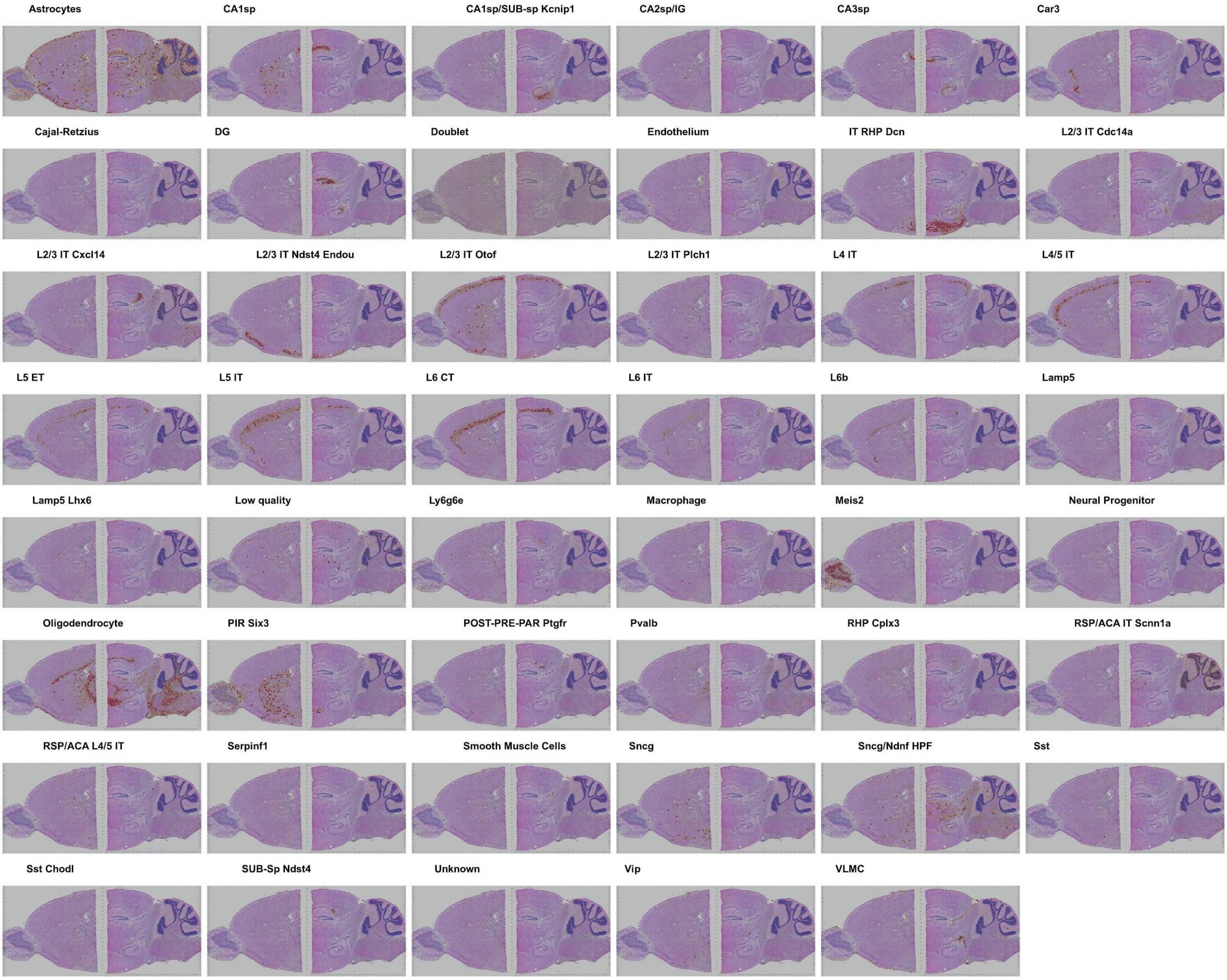


**Supplementary Figure 5 |** Predicted cell-type proportion within each capture location replicate 2.

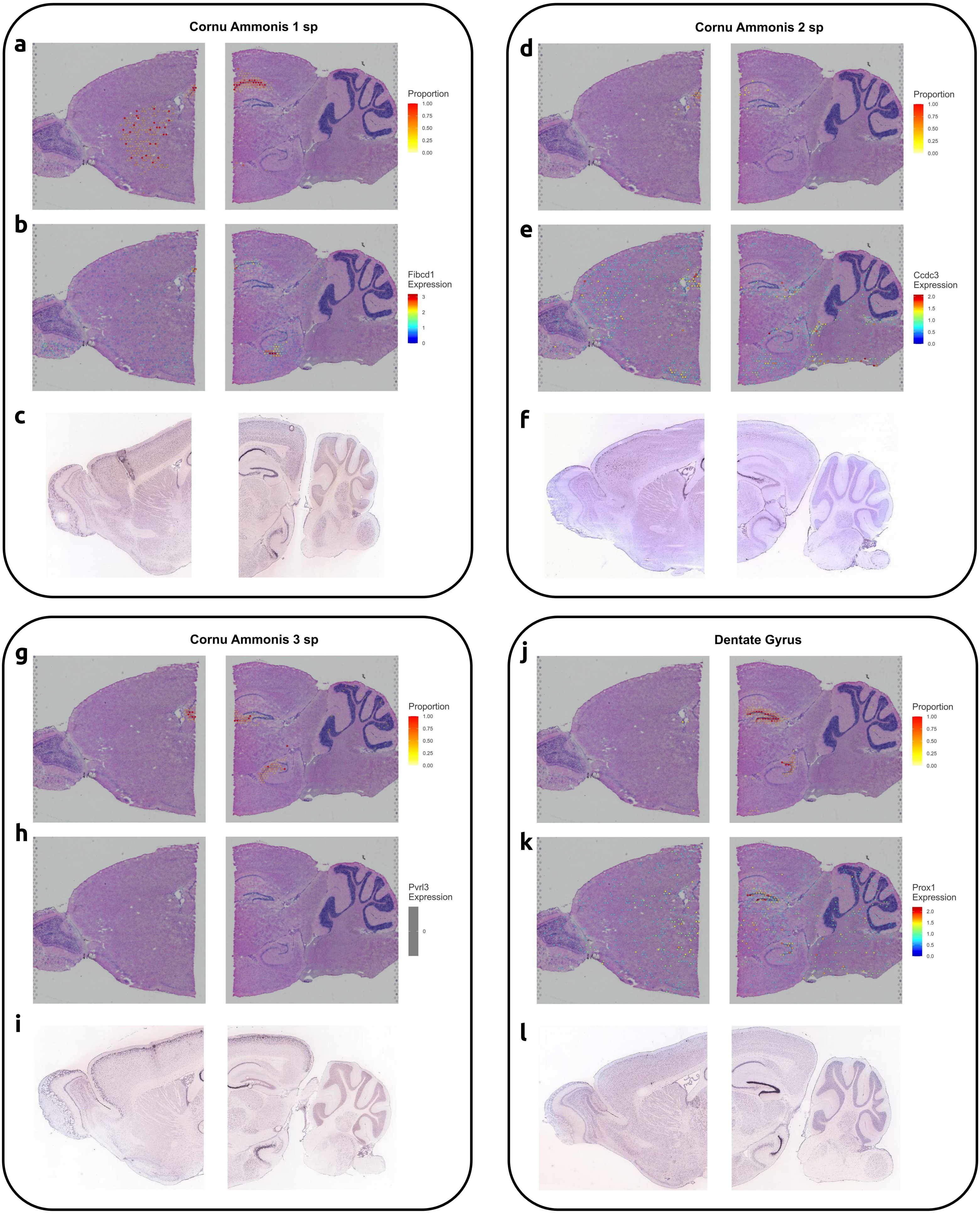


**Supplementary Figure 6 | Hippocampal cell type mapping on sagittal adult mouse brain anterior and posterior slices. a, d, g, j**, Proportions of the hippocampal cells from the reference atlas within capture locations. **b, e, h, k**, Canonical marker gene expression in each capture location. **c, f, i, l**, Reference In Situ Hybridization labelling of the canonical marker genes from the Allen Institute to validate the cell types are predicted on their correspondent structure.

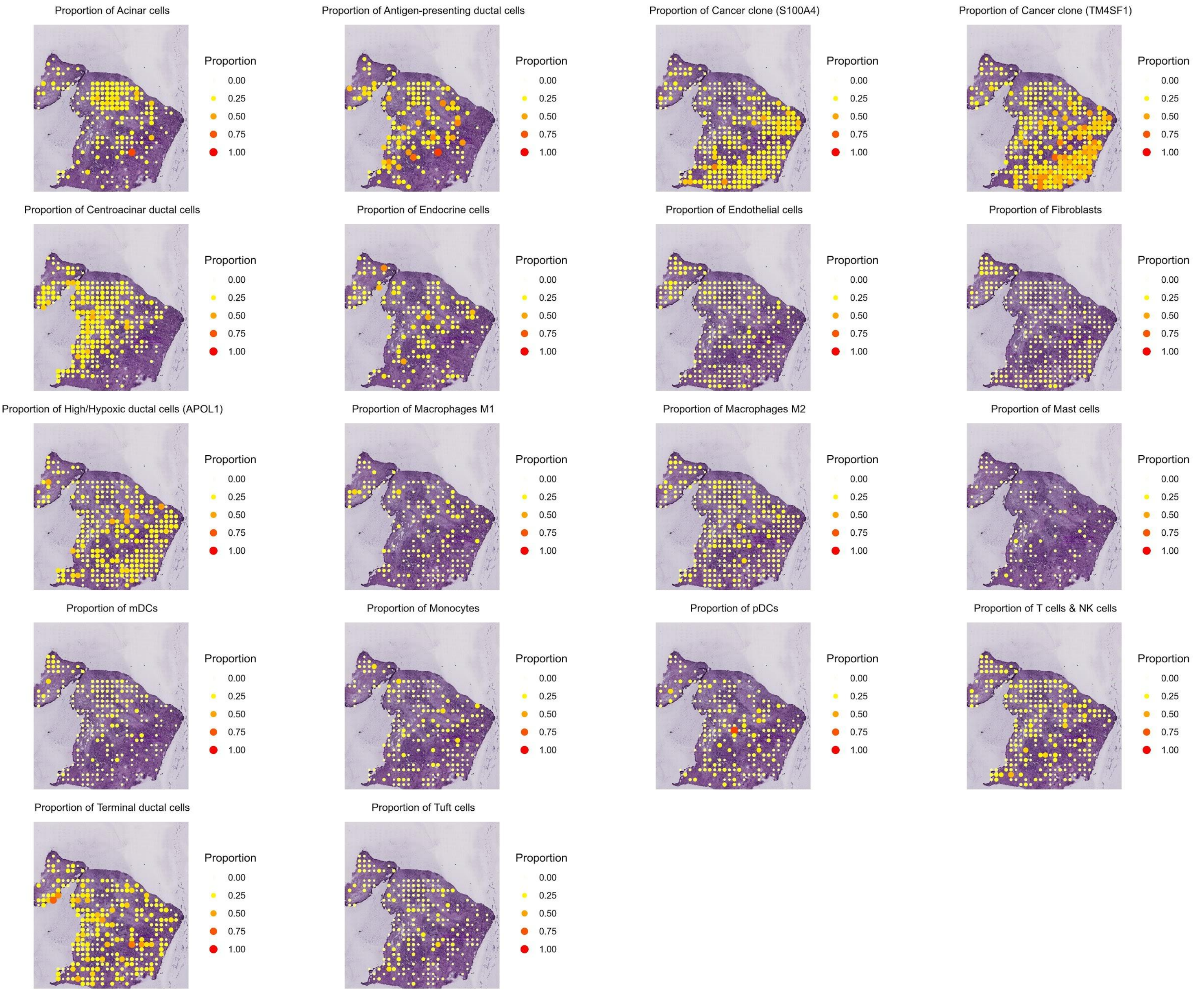


**Supplementary Figure 7 |** Predicted proportion within each capture location of the cell types in the paired PDAC-A indrop dataset on the PDAC-A slide.

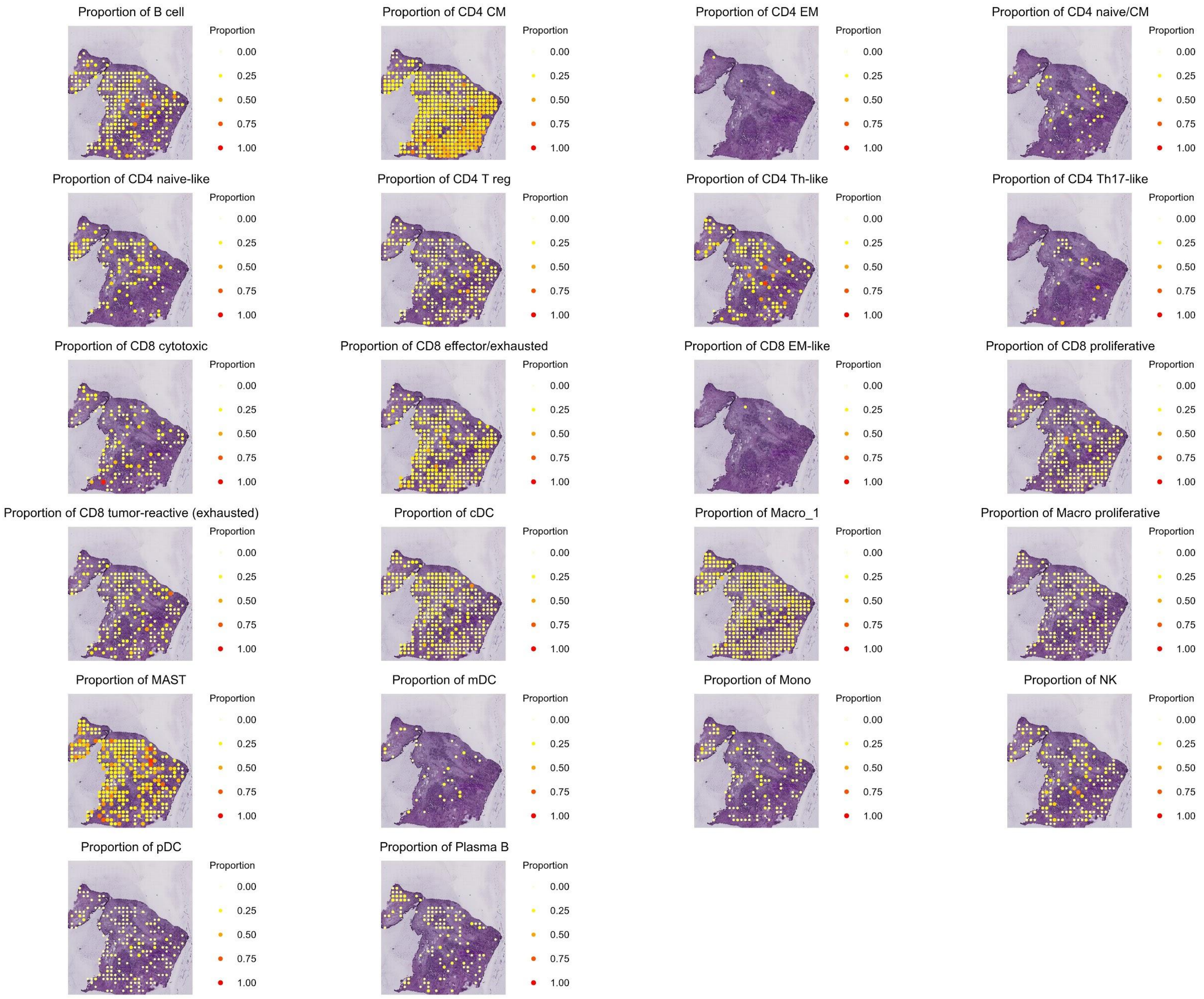


**Supplementary Figure 8 |** Predicted proportion within each capture location of the cell types from the immune reference dataset on the PDAC-A slide.

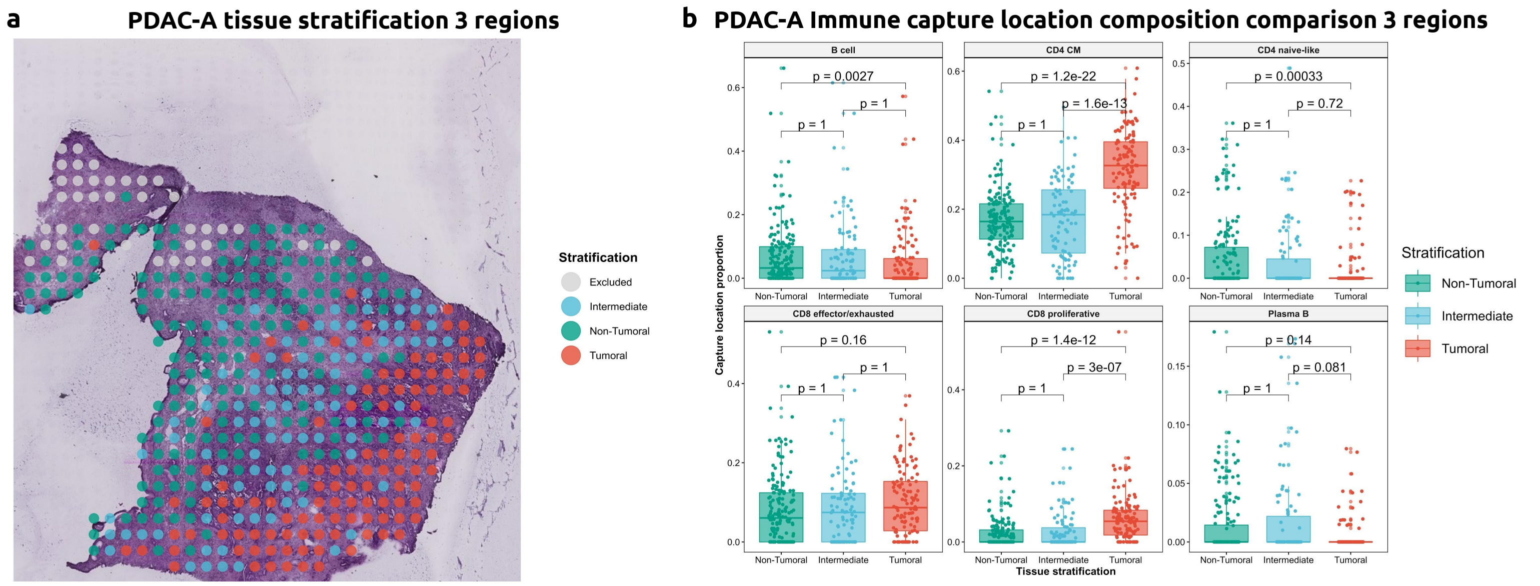


**Supplementary Figure 9 |** Predicted proportion within each capture location of the cell types in the paired PDAC-B indrop dataset on the PDAC-B slide.

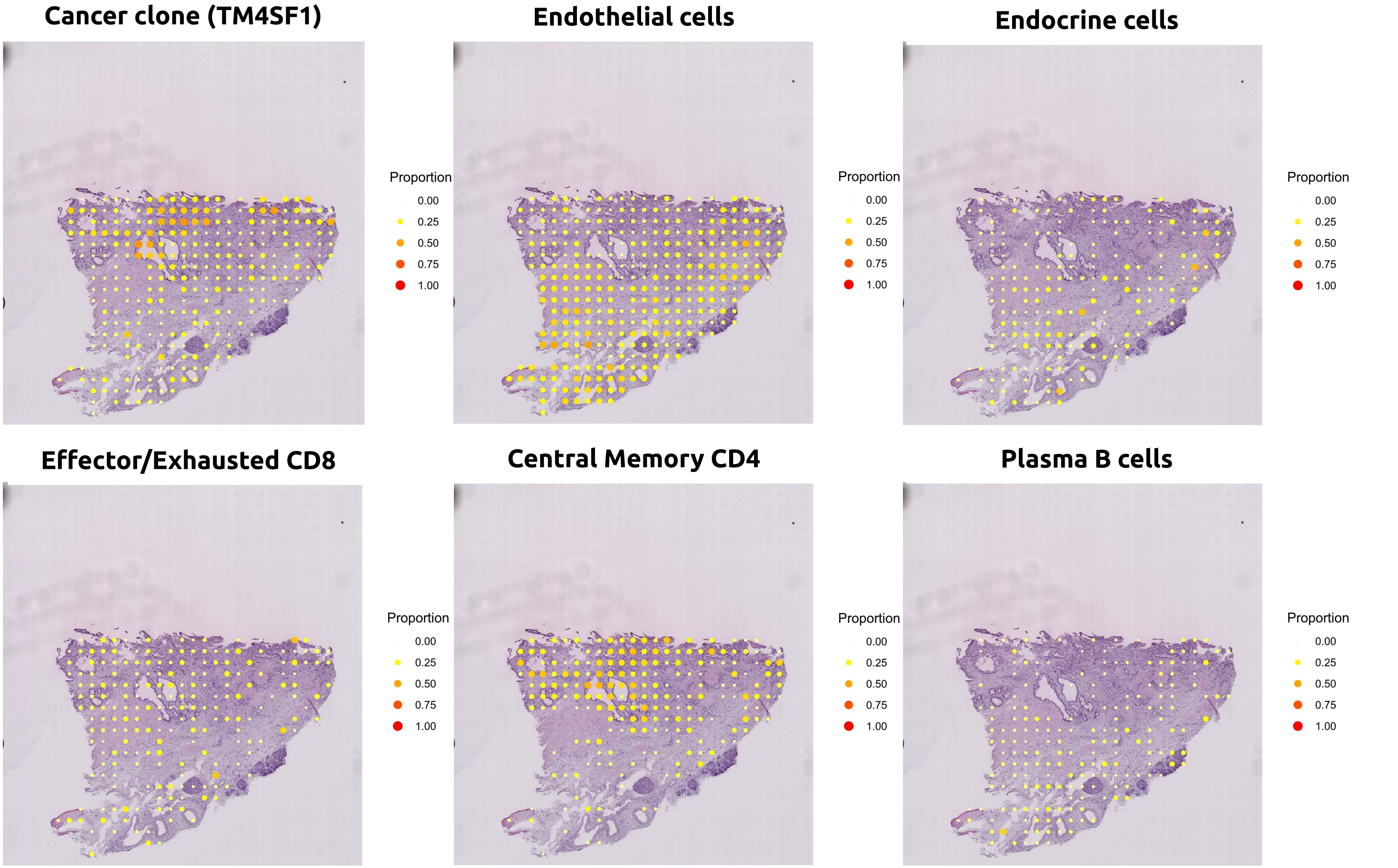


**Supplementary Figure 10 | Tissue stratification and cell type proportion comparison within the spots by region. a**, Tissue stratification by tumoral, intermediate,and non-tumoral capture locations. **b**, Cell type proportion comparison within each spot between tumoral, intermediate, and non-tumoral sections. Central memory CD4 and proliferative CD8 cells are found at higher proportions within the tumoral region. Naive CD4 and B cells show higher proportions in the non-tumoral section.

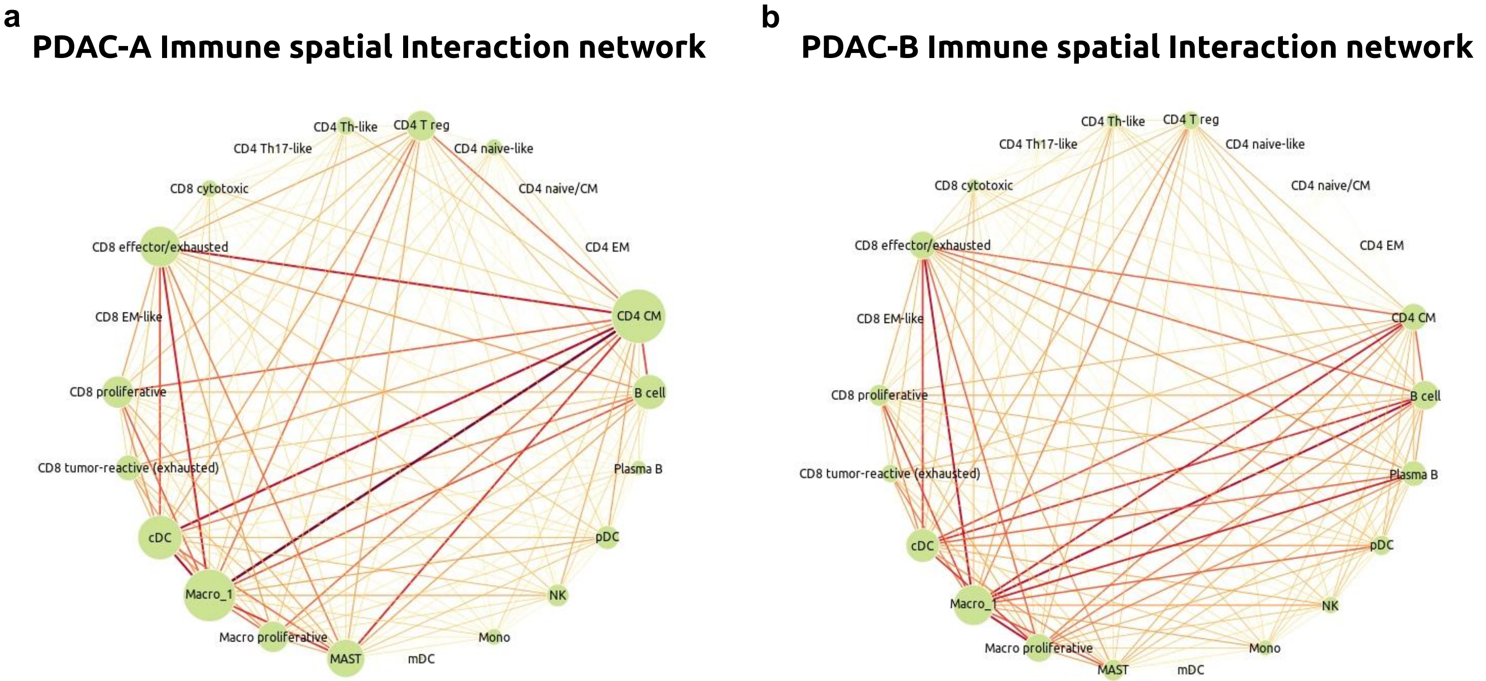


**Supplementary Figure 11 |** Spatial interaction network representing the frequency with which cell types co-localized together. **a**, Immune cell network for PDAC-A. **b**, Immune cell network for PDAC-B

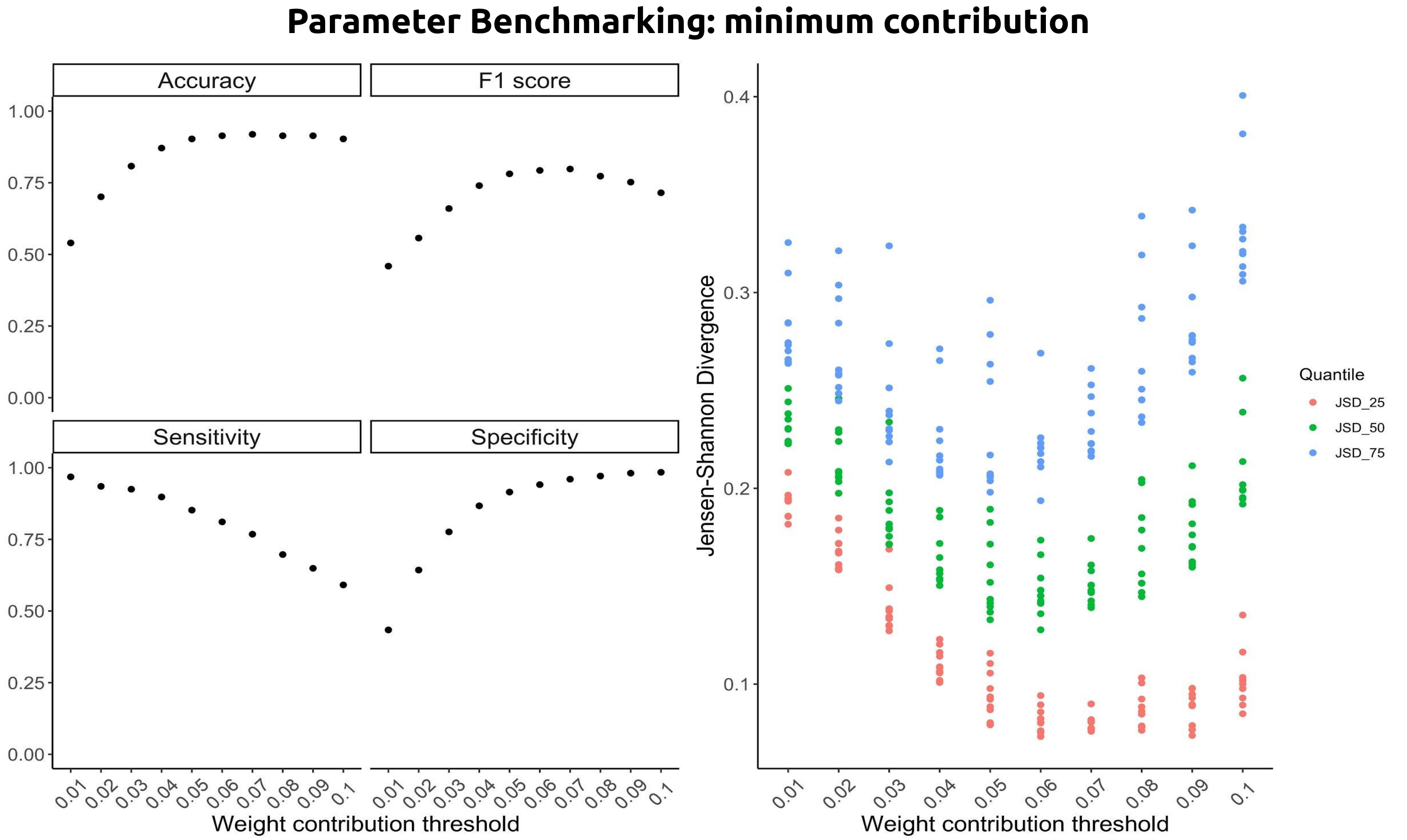


**Supplementary Figure 12 |** Optimizing threshold parameter to select minimum contributing weight to consider a cell type present within a spot.
